# Results from a conservation initiative for parrotfishes in the Colombian Caribbean

**DOI:** 10.1101/2021.09.27.462015

**Authors:** Juliana López-Angarita, María del Pilar Restrepo Orjuela, Katherine Guzmán Peña, Dairo Escobar

## Abstract

Parrotfish (Family Scaridae) are a family of herbivorous fishes crucial to coral reef health, particularly for Caribbean reefs due to their declining coral cover. However, despite parrotfish are fully protected in some countries, they are still heavy fished in most of their Caribbean range. The consequences of this targeted fishery in the Colombian Caribbean are not fully understood due to a lack of local conservation and management resources. This research aimed to evaluate and enhance the conservation status and protection of parrotfish among local communities in the National Natural Park *Corales del Rosario and San Bernardo*. Underwater visual census surveys (UVC) were undertaken to evaluate reef fish community structure, and participatory education campaigns and activities were carried out with local communities to raise awareness about parrotfish ecology and their functional role in conserving Caribbean coral reef ecosystems of Colombia. UVC showed parrotfish to be dominant in the fish community, yet there was evidence of exploitation of large adults by selective fishing. Conflicts exist between the community and environmental authorities because fishing regulations are not clear, and the level of enforcement is insufficient. Parrotfish are sold to tourists, as ‘red snapper’ to fulfil high seafood demand since commercially valuable fish are now scarce. However, following intensive awareness-raising activities developed as part of this study, the community has started to recognize the vital ecological role of parrotfish in coral reef systems, and are suggesting a redrafting of fishing legislation by the environmental authorities, in order to recognise and incorporate the traditional fishing rights of human communities living within the MPA. Lobbying for the protection of parrotfish and inclusion of local communities in decision-making will take time, but this research represents the crucial first steps towards sustainable practice and cooperative alliances in the Colombian Caribbean.

## 1. Introduction

The Caribbean has suffered region-wide declines in coral reef health since the 1970’s (Gardner et al., 2003). According to the Atlantic and Gulf Rapid Reef Assessment (AGRRA) Program, who have been conducting regional surveys of coral reef health for 20 years, the regional average of macroalgal cover is 40%, while coral cover currently stands at 13% (AGRRA, 2018). Concomitant to this degradation is the intensification of fishing pressure that has caused overfishing of commercial and herbivorous species in most locations (Shantz et al. 2020). This is a worrying situation for many small island developing states and small coastal communities that depend strongly on small-scale fishing for their livelihoods while relying on many other ecosystem services provided by reefs. The sustainable management of reef resources is a complex interdisciplinary process requiring the evaluation and integration of multiple factors (biological, ecological, socioeconomic and institutional) to understand the various dynamics affecting the ecosystem (Pollnac et al. 2010).

Parrotfish are herbivores that fulfil a key role in coral reef health by consuming macroalgae, which inhibits coral growth by dominating bare substrate and restraining coral recruitment (Hughes et al. 2007; Ledlie et al. 2007; Mumby et al. 2007). By scraping algae from rock and coral, parrotfish also act as a major bioeroder, producing large amounts of sand as faeces (Morgan & Kench, 2016). Fishing targets large individuals and as such can shift the size structure of populations, increasing the density of small individuals, which are less efficient in controlling algae (Durán & Claro 2008). Shantz et al. (2020) showed that algal cover of Caribbean coral reef communities was negatively correlated with the density of large parrotfish. Therefore, the depletion of parrotfish populations may cause rapid and dramatic changes in reefs structure and functionality, especially when combined with eutrophication, potentially leading to a phase shift from a coral-dominated to an algae-dominated state (Bellwood et al. 2004; Fabricius 2005; Mumby et al. 2007). For this reason, parrotfish are widely acknowledged as important enhancers of coral reef resilience (Hughes et al. 2007).

In this study we assess the abundance and structure of parrotfish populations in the National Natural Park *Corales del Rosario and San Bernardo*. Additionally, we determine the scale and characteristics of the parrotfish fishery by gathering socioeconomic indicators to assess the impact of parrotfish extraction on reef health and future provision of fishing products. The project also focused on increasing community awareness about the important functional role of parrotfish in coral reefs by conducting educational activities and protection campaigns.

## 2. Methods

### 2.1. Study site

The National Natural Park *Corales del Rosario and San Bernardo* (CRSB) located in the Colombian Caribbean comprises two archipelagos, 30 small islands, and coastal lagoons harbouring mangrove forest, tropical dry forest, rocky shores, seagrass beds and coral reef within a total area of 120,000 Ha. Established in 1977 as the first marine park in the country, CRSB has an important conservation value protecting the most extensive, diverse and developed coral reefs of the continental shelf of Colombia (Pineda et al. 2006). Most of the islands within the CRSB do not belong to the protected area and have been inhabited by native Afro-Colombian communities (ethnical minority) from about 300 years (Durán 2009). Since its inception, the park has suffered a series of conflicts between local communities, government & environmental authority, involving land tenure issues, rules compliance, and local participation in management (Durán 2006).

Due to its proximity to the city of Cartagena and a dramatic increase in tourism, CRSB is the most visited national park in Colombia (Pineda et al. 2006). Coastal development fostered by tourism has led to overexploitation of marine resources, where fishing has become one of the main threats to coral reef health since it must meet the increasing demand of the tourism industry as well as sustaining the resident population (Pineda et al. 2006). Coral bleaching in recent years has caused the mortality of nearly 95% of Elkhorn coral (*Acropora spp*.) and other species (E. Zarza, *pers. comm*.), and water pollution from regional sewage and riverine contaminants further inhibit coral resilience and recovery (Bejarano et al., 2016). A recent increase of fishing pressure of commercially valuable piscivorous fish species such as snappers and barracudas in the region (Martínez-Viloria et al. 2011) has led to a system dominated by herbivorous parrotfish (Scaridae) that traditionally have had no commercial value, but are now targeted.

### 2.2. Underwater visual surveys

To determine the status of fish populations we conducted underwater surveys between 2008 and 2009 at 10 sites located inside and 6 sites outside CRSB limits. We sampled areas in shallow (3–15 m) highly developed reefs, ensuring geomorphological and environmental similarities. Underwater visual surveys were conducted by swimming along 2 × 50 m belt transects (100m^2^) recording all individual fish of parrotfish family and other important families (Acanthuridae, Lutjanidae, Serranidae, Haemulidae, Carangidae, Scombridae, Sphyraenidae) and estimating total fish length. Between 4 and 10 transects were carried out for each site. Fish biomass was calculated by converting length estimates to weight using length-weight conversion equation: *W= aTL*^*b*^. All fitting parameters were obtained from FishBase (www.fishbase.org). We calculated herbivore biomass, total fish density and biodiversity for each site and then compared these parameters between MPA and non-MPA surveys using Wilcoxon Rank Test for non-parametric data and ANOVA for parametric data.

### 2.3. Community surveys

To characterize fishing activity and the parrotfish market we i) conducted household surveys and participatory workshops with local communities to explore social context (implementing the following tools: productive profile, historic graph, seasonal analysis, and problems & solutions tree); ii) performed surveys of fishermen about their knowledge of the ecological role of parrotfish within the coral reefs; iii) conducted surveys of tourists visiting the National Park to determine their knowledge of these species and their willingness to pay for consuming parrotfish; and iv) performed surveys of restaurant managers inside the Park to establish the gastronomic demand of tourists.

### 2.4. Awareness raising activities

We worked closely with the CRSB environmental education staff who helped us to develop awareness raising activities. We organized diverse activities and adapted the strategies according to the culture of the region to make the message more understandable and valuable for the local community. We organised: i) educational workshops and talks with fishermen; ii) “the parrotfish day”; iii) a song “la champeta del loro”; iv) art workshops and talks in local schools (held in the communities of Isleta, Isla Grande, Múcura y Santa Cruz El Islote); v) a theatrical play with the local school in Isla Grande; vi) sport events (football for adults, softball for kids); vii) educational campaign for tourists by printing booklets and posters with key information about parrotfish conservation and distribute them in key sites of Cartagena city and the park (tourism centres, ports, diving shops, hotels, restaurants, eco-hotels); viii) talks during high season in tourism boats; ix) workshops about the biology and ecology of parrotfish, school snorkelling activities with eco-guides, and designed underwater guides & posters for parrotfish identification.

## 3. Results

### 3.1. Underwater visual surveys

Parrotfish (Scaridae) were the most important family comprising 48% of the total number of fish, followed by surgeonfish (Acanthuridae) 24%, and Haemulidae 17% (Figure 2). Diversity indices showed slightly higher values from inside the MPA, but values were not statistically significant from unmanaged sites (Table 1). Fish density inside and outside were non significantly different (Wilcoxon P>0.05) (Figure 2). Herbivore biomass (mean grams ±SD) inside the MPA (32.15 gr ± 12.14) was slightly higher than in populations outside the MPA (23.80 gr ± 11.34) however differences were not statistically significant (ANOVA P>0.05).

**Figure 1.**
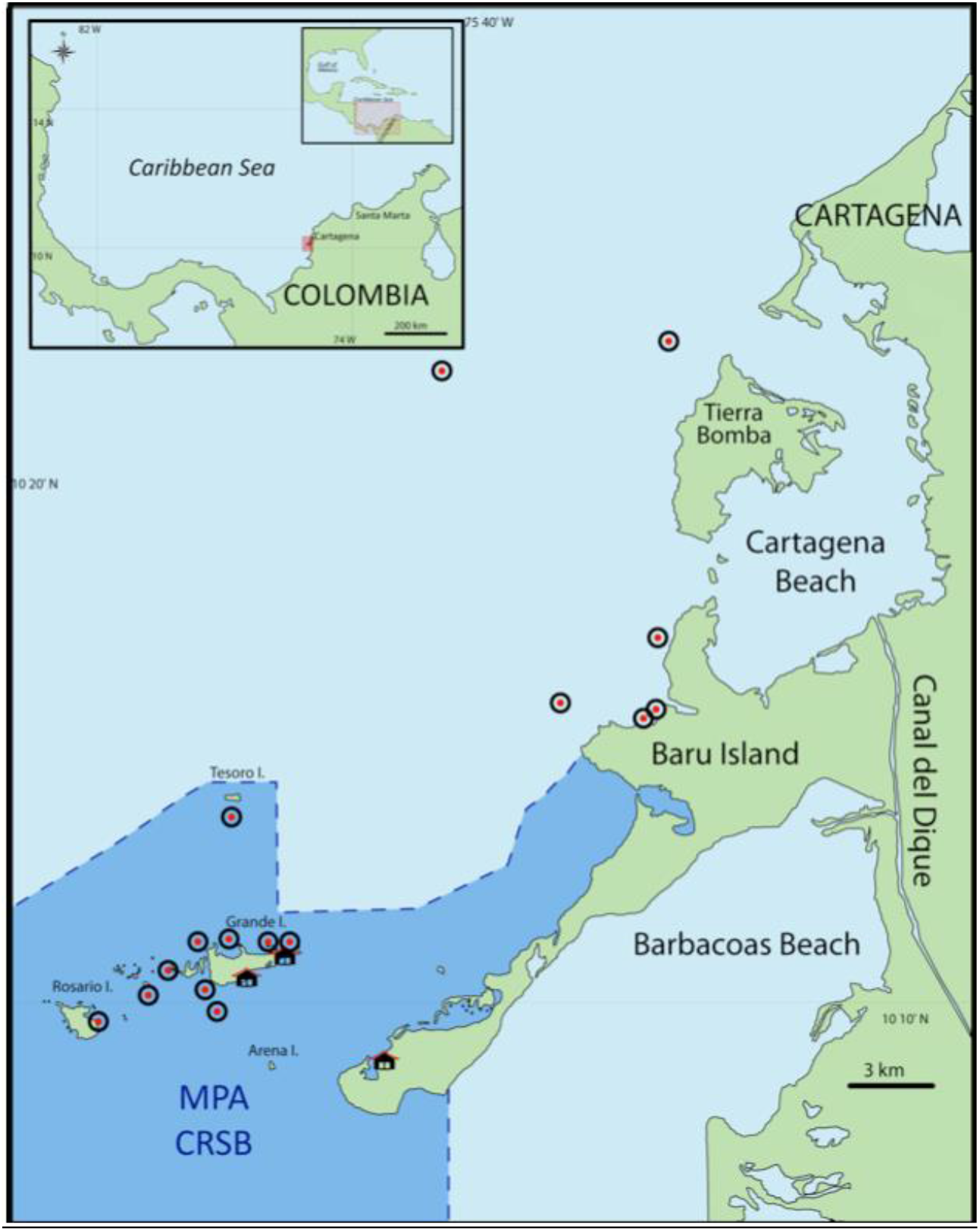
Study area with biophysical sampling sites inside and outside the MPA are shown with a circle; and local communities with a house icon.

**Figure 2.**
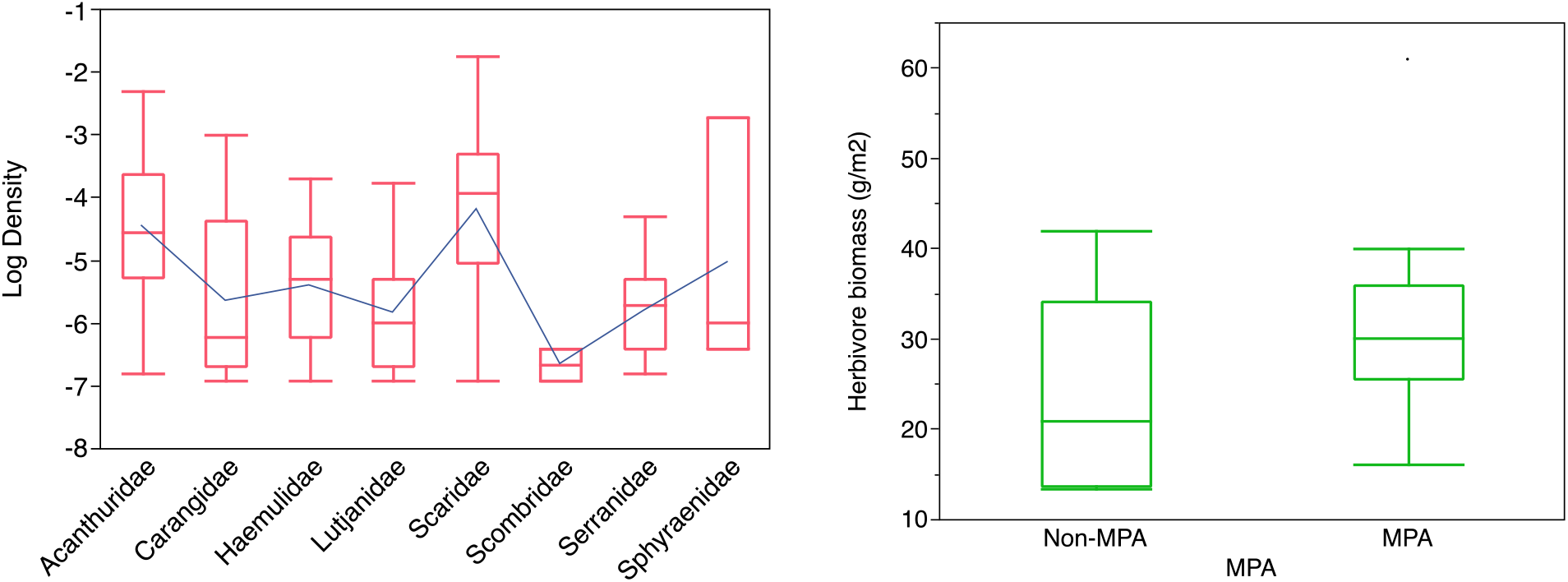
Density (#/m^2^) of all fish families surveyed (left panel). Herbivores such as parrotfish (Scaridae) and surgeonfish (Acanthuridae) are more abundant per square meter than all the other commercial families. Mean biomass of herbivore populations inside and outside the MPA. Biomass is higher inside but differences are not significant (right panel). Blue line is a visual guide only.

**Table 1.** Diversity indexes for fish populations in sites inside and outside CRSB and significance (p) obtained from ANOVA comparisons.

| Diversity Index | MPA | Outside MPA | P |
| --- | --- | --- | --- |
| Shannon index (H') | 2.56 | 2.48 | 0.521 |
| Rarefied richness | 32.5 | 30 | 0.857 |
| Evenness (J') | 0.72 | 0.73 | 0.378 |

Parrotfish density was significantly different between species (Wilcoxon P<0.0001). The most abundant species were: striped (*Scarus croisensis*), redband (*Sparisoma aurofrenatum*), and stoplight parrotfish (*Sparisoma viride*). Midnight parrotfish (*Scarus coelestinus)* was the least common species (1 individual outside the MPA), and blue *(Scarus coelurus) &* rainbow parrotfish *(Scarus guacamaia)* were not observed during surveys (Figure 3).

**Figure 3.**
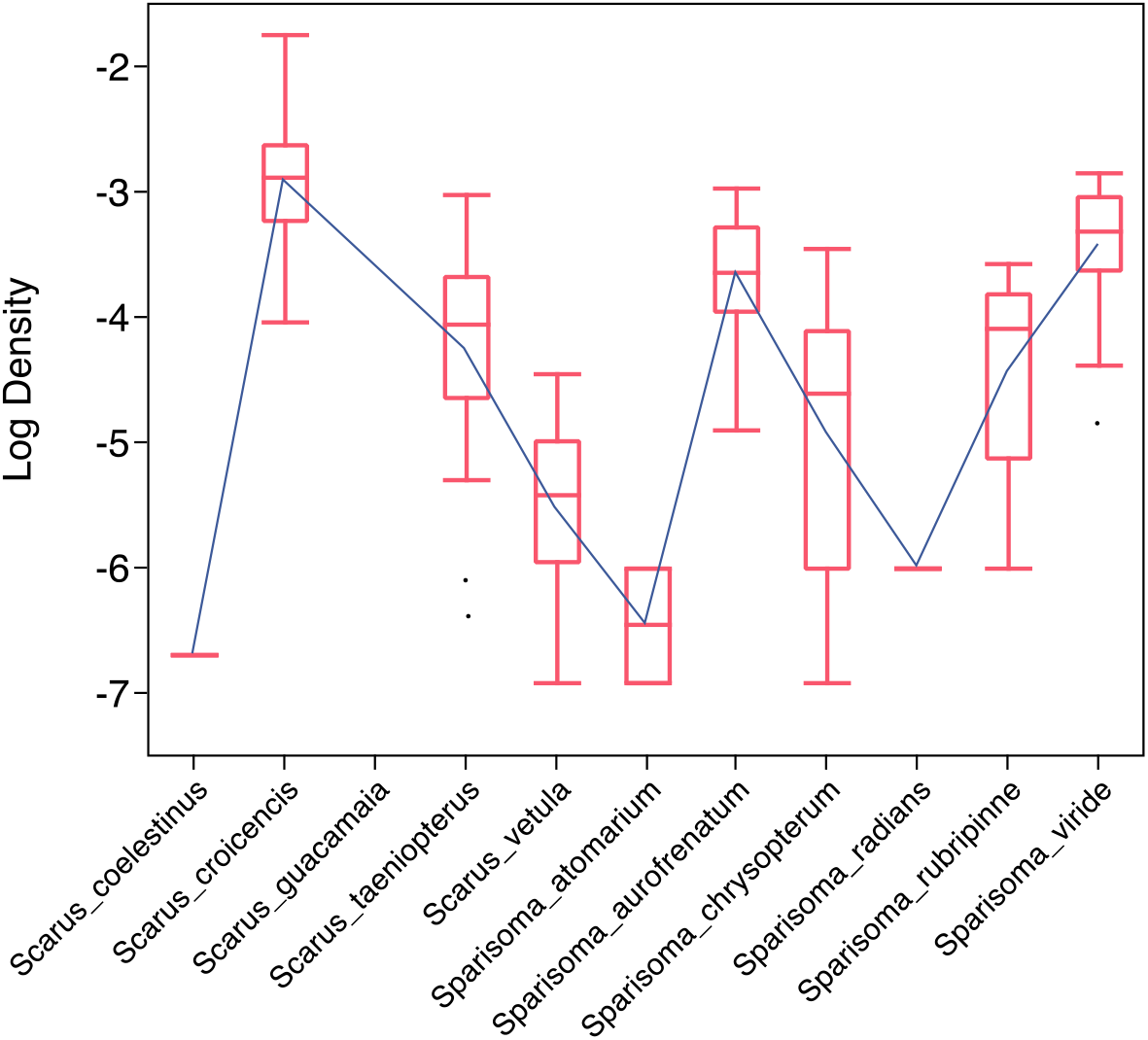
Density per m^2^ of parrotfish species (Scaridae), there were significant differences between them, showing the common and least common species in the ecosystem. Blue line is a visual guide only.

Mean and maximum size data (total length) for observed species were compared between documented values in FishBase (“common length” and “Max length”) in order to calculate percentage difference between observed sizes and those documented from elsewhere in the Caribbean (Table 2). Mean parrotfish size observed in CRSB was 12.69% smaller inside and 18.68% smaller outside the park in comparison with documented mean sizes. The only case where the observed size was higher than the documented was *S. chrysopterum*, showing larger mean size by 0.2cm inside CRSB. Stoplight parrotfish showed the largest size reduction with an alarming value of ∼30% less than documented values, with most individuals exhibiting sizes between 15-30 cm (Figure 4).

**Table 2.** Table of comparison between documented and observed mean & maximum sizes (total length) of adult parrotfish. Sizes are in centimetres and means are listed with the standard deviation (± SD). Documented values were obtained from FishBase (www.fishbase.org).

| Species | Documented values |  | Non MPA |  |  |  | MPA |  |  |  |
| --- | --- | --- | --- | --- | --- | --- | --- | --- | --- | --- |
|  | Mean size | Max size | Mean size observed | Mean size difference | Max size observed | Max size difference | Mean size observed | Mean size difference | Max size observed | Max size difference |
| *Scarus croicensis* | 18.0 | 35.0 | 15.5 ± 3.5 | 14.1% | 25.0 | 28.57% | 17.5 ± 4.1 | 3.01% | 35.0 | 0.00% |
| *Sparisoma aurofrenatum* | 20.0 | 28.0 | 16.2 ± 3.7 | 18.8% | 25.0 | 10.71% | 17.7 ± 4.2 | 11.60% | 25.0 | 10.71% |
| *Scarus taeniopterus* | 22.0 | 35.0 | 18.7 ± 4.0 | 14.8% | 25.0 | 28.57% | 19.3 ± 5.4 | 12.35% | 35.0 | 0.00% |
| *Sparisoma chrysopteron* | 25.0 | 46.0 | 22.6 ± 4.7 | 9.6% | 30.0 | 34.78% | 25.2 ± 7.4 | -0.87% | 45.0 | 2.17% |
| *Sparisoma rubripinne* | 30.0 | 47.8 | 22.9 ± 5.2 | 23.6% | 30.0 | 37.24% | 26.8 ± 7.1 | 10.69% | 45.0 | 5.86% |
| *Scarus vetula* | 32.0 | 61.0 | 27.0 ± 3.7 | 15.5% | 30.0 | 50.82% | 25.5 ± 7.7 | 20.20% | 45.0 | 26.23% |
| *Sparisoma viride* | 38.0 | 64.0 | 25.8 ± 7.5 | 32.2% | 40.0 | 37.50% | 25.9 ± 7.0 | 31.87% | 45.0 | 29.69% |

**Figure 4.**
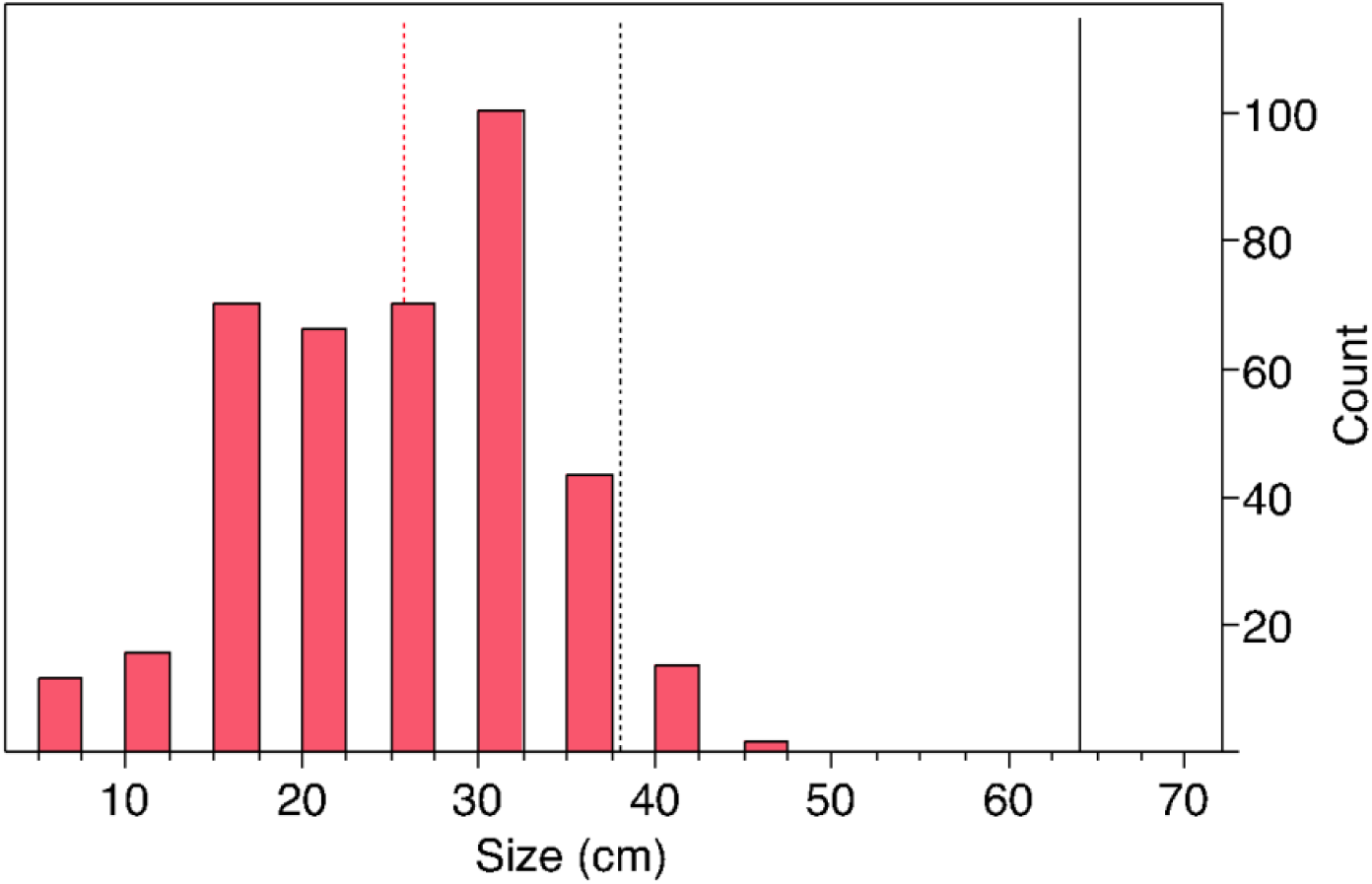
Frequency distribution of stoplight parrotfish (Sparisoma viride) population sizes. Solid black line represents the maximum documented size for this species, dotted black line represents the mean size documented and dotted red line shows the mean size observed in this study.

The mean abundance of parrotfish per count (transect) inside and outside CRSB was calculated. In general smaller species were more abundant than bigger species. However, *S. radians, S. atomarium* and *S. viride* were the exception to this trend, because the first occurs generally in seagrass beds, the second is hard to record accurately due to crypsis (Hawkins & Roberts 2003), and the last seems to be highly resilient to fishing pressure. Counts outside the CRSB had a higher proportion of small sized species. All the parrotfish species present in the Caribbean were seen during the surveys except for the largest *species, S. coelurus and S. guacamaia* (Figure 5).

**Figure 5.**
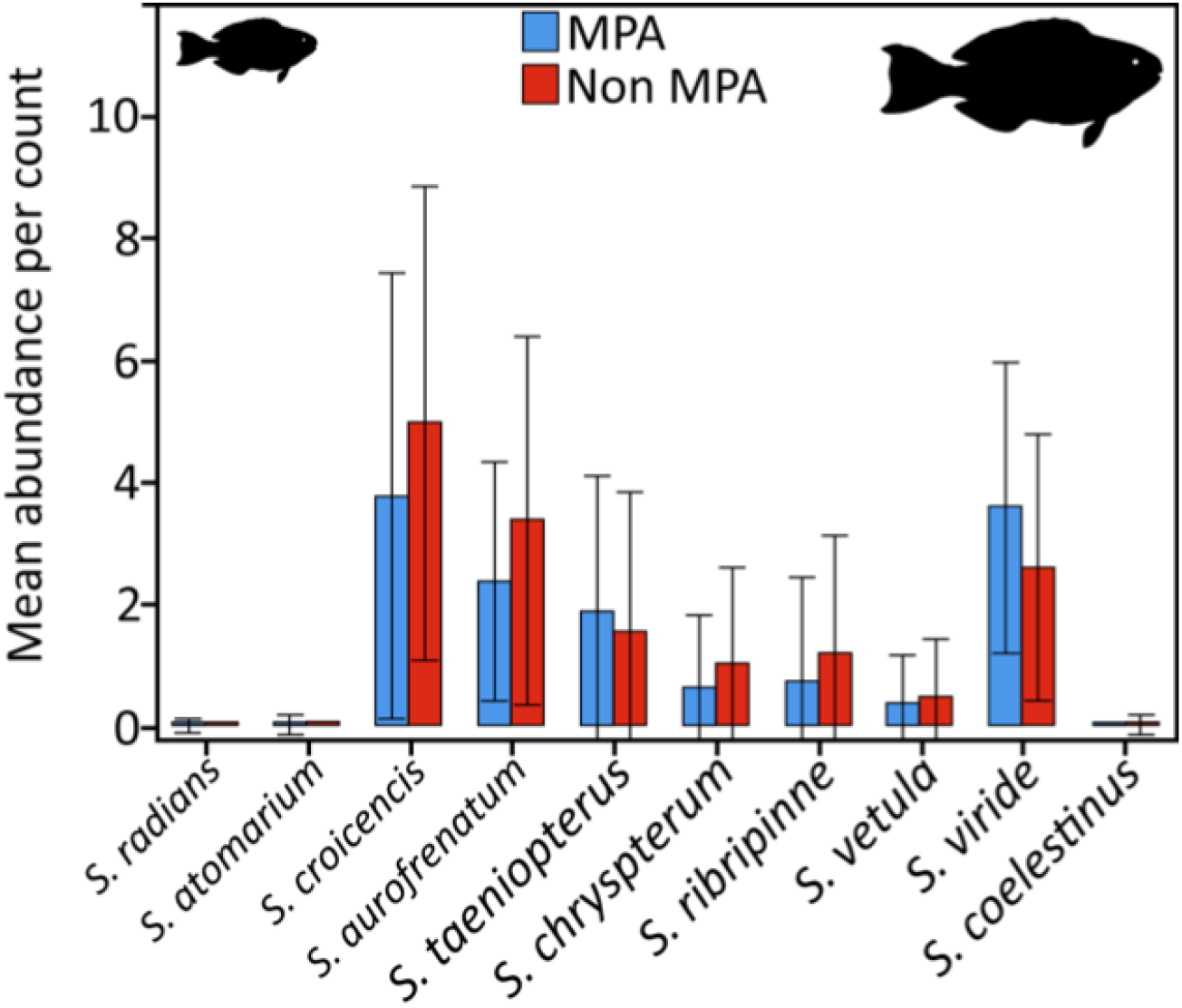
Mean abundance of parrotfish per count. Species are arranged along the x-axis in order of increasing documented size and abundance for counts inside and outside CRSB are shown.

### 3.2. Community surveys

Sixty-eight household surveys were carried out in the biggest communities of Rosario (15 in Isleta, 35 in Isla Grande and 18 in Barú). Surveys showed that fishing is the principal economic activity of the majority of households and that generates an average income of US $156 per month. Including other economic activities, the average income of a household per month is US $189, slightly over half the national minimum wage ($318 per month). Additionally, 96% of head of households affirmed that fishing resources have diminished in the last 10 years and 91% proposed that management of resources should be made through agreements between community and authorities.

Only 12% of the 39 fishers surveyed in Rosario (Isleta, Isla Grande and Barú) reported to target parrotfish. These fishers estimated that parrotfish represents 15% of the total daily capture. When asked about the use of parrotfish, 88% confirmed it is used for household consumption. On the other hand, during fisher group workshops, the historical analysis showed that the parrotfish market began around 2000 for commercial purposes (Table 3).

**Table 3.** Historic graph showing how important variables related to parrotfish evolved in time.

| Variable | Before 1980 | 1980-1990 | 1990-2000 | 2000-2005 | 2005-2010 |
| --- | --- | --- | --- | --- | --- |
| Parrotfish extraction | Household consumption around 1985 because scarcity of other species | Low extraction | Low extraction | Extraction increases for commercialization | Very high extraction for commercialization |
| Parrotfish abundance trends | Always abundant |  |  |  |  |
| Fishing gears |  | Pole spear | Pole spear | Pole spear | Pole spear |

Our observations in the field and discussions with stakeholders confirm that fishermen catch parrotfish to satisfy the gastronomic demand from tourists, where parrotfish are sold disguised as snapper in the typical Caribbean lunch (whole fried fish, fried plantains, coconut rice and salad). Fishers sell one individual parrotfish to restaurants for US $0.50, who then will sell it to tourists at the price of snapper ($6 plate) (Table 4). This entire process is conducted in a confidential manner as fishermen are aware restaurants trick tourists and because park staff have recommended fishers against capturing parrotfish.

**Table 4.**
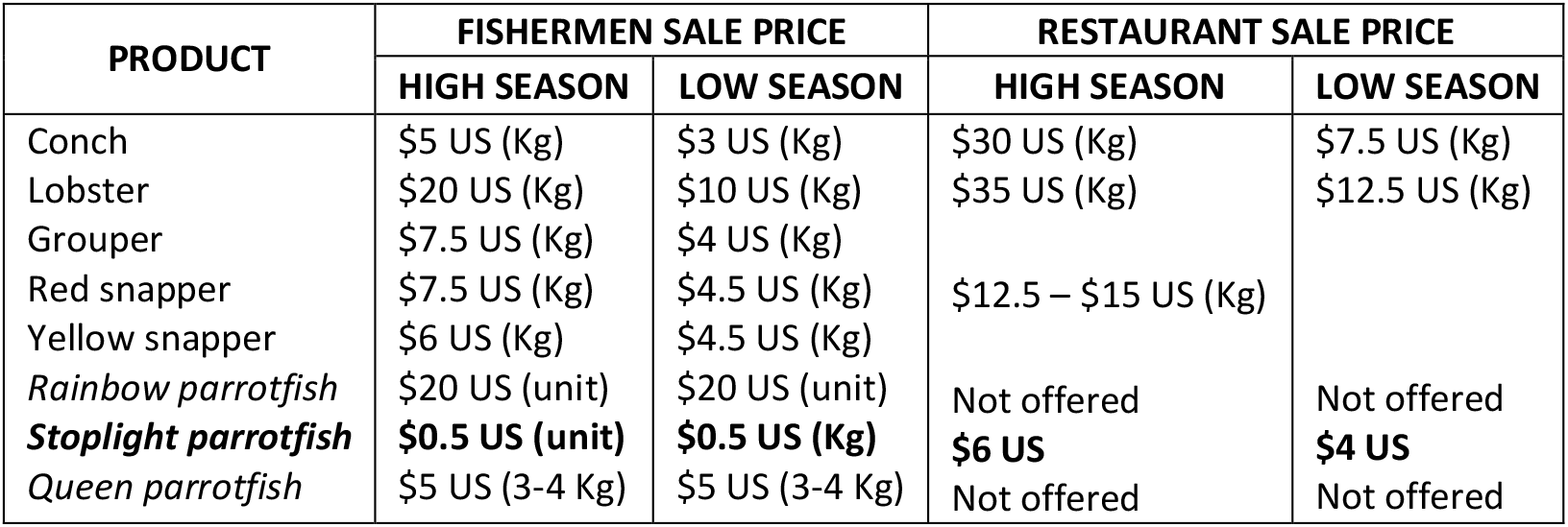
Information of the market of commercial and herbivore species for the community of Isla Grande obtained during the workshop.

| PRODUCT | FISHERMEN SALE PRICE |  | RESTAURANT SALE PRICE |  |
| --- | --- | --- | --- | --- |
|  | HIGH SEASON | LOW SEASON | HIGH SEASON | LOW SEASON |
| Conch | \$5 US (Kg) | \$3 US (Kg) | \$30 US (Kg) | \$7.5 US (Kg) |
| Lobster | \$20 US (Kg) | \$10 US (Kg) | \$35 US (Kg) | \$12.5 US (Kg) |
| Grouper | \$7.5 US (Kg) | \$4 US (Kg) | | |
| Red snapper | \$7.5 US (Kg) | \$4.5 US (Kg) | \$12.5 – \$15 US (Kg) | |
| Yellow snapper | \$6 US (Kg) | \$4.5 US (Kg) | | |
| Rainbow parrotfish | \$20 US (unit) | \$20 US (unit) | Not offered | Not offered |
| **Stoplight parrotfish** | **\$0.5 US (unit)** | **\$0.5 US (Kg)** | **\$6 US** | **\$4 US** |
| Queen parrotfish | \$5 US (3-4 Kg) | \$5 US (3-4 Kg) | Not offered | Not offered |

In restaurant surveys (n=6), only 16% of establishments openly admitted to sell parrotfish on their menus, yet when asked if they would buy parrotfish offered by fishers, 50% of the restaurant owners were prepared to buy these species and offer it in their restaurants as according to them, it is cheaper and tastes good. Half of restaurants declared that few tourists are capable of identifying the species by its colour, shape or size. We interviewed the families that manage 6 restaurants in Isla Múcura and they confirmed that each restaurant serves 15-20 parrotfish lunches in low season, this number increases to 120-200 in high season, which translates into 720-1800 parrotfish served per day in the island.

We conducted 109 surveys to tourists in Cartagena and inside CRSB. 34% of tourists had heard about the parrotfish, of which only 13.5% knew their ecological importance. Once informed of the importance of this fish to coral reef health, tourists were asked if they were willing to consume parrotfish at the same price as other commercial species, to which 89% answered negatively, and 80.7% informed they would not consume it even if it was half the price.

### 3.3. Awareness raising activities

In the educational workshops we used didactic tools to evaluate the ecological knowledge of fishermen. Many fishermen had misconceptions about the role of parrotfish, and most were unaware of its importance. With the help of local school children and park officers we organized a “Parrotfish Day” for the community of Isla Grande. The event included a theatrical play of the ecological role of the parrotfish, a pictographic exposition with the best selection of children’s drawings of the parrotfishes’ role in the ecosystem, and a live concert featuring the famous local musician Charles King. The concert represented the first live performance of “*La champeta del loro*” (https://youtu.be/J4nl8jPbkUU), a parrotfish conservation song written and recorded for the project and performed for the whole community. Additionally, we organized a “parrotfish sports week” with adults and children, awarding prizes from the project (hats, uniforms, notebooks, CD’s).

For the awareness raising campaign for tourists we gave informative talks during the high season to 500 people and we trained 7 National Park volunteers to be conservation educators (in charge of giving information to tourists at the port prior to departure to the islands). Posters were placed in strategic sites, creating an agreement with the owners/managers of the establishments to post it permanently and disseminate information. We estimate that a total of 6500 people were influenced by our tourist campaign. In Isla Grande we trained the leaders of the native eco-guides group with a workshop, snorkelling activities and parrotfish identification guides, and posted two permanent posters on the island that will serve as stop stations to talk about parrotfish when they are giving the tour around the island with tourists.

Once the project was finalised, we surveyed 39 fishermen and 8 park officers to determine the effectiveness of our raising awareness activities. 35% of fishermen admitted to having learned about the importance of parrotfish from the CLP project; 100% expressed the importance of conserving them; 89% were willing to stop fishing parrotfish; and 77% proposed to prohibit parrotfish capture as a method to protect it. Regarding park officers, 87.5% understood the ecological role of parrotfish, and 50% of them admitted to having learned this information from the educational activities related to the project.

## 4. Discussion

Statistical analysis showed that fish density, biodiversity and biomass inside the MPA were not significantly different from reefs outside the MPA. This suggests that fishing pressure and its impacts on fish population is similar inside and outside the reserve, despite the existence of fishing regulations in the MPA. Parrotfish were the most representative family comprising 48% of the fish, however densities between parrotfish species were significantly different. Large size species were the least common or totally absent in surveys (e.g. *S. guacamaia* listed as vulnerable on the IUCN red list), whereas smaller sized species were more abundant in the communities. There is an evident reduction in total length for all the species in CRSB compared with documented common and maximum sizes, suggesting the effects of overfishing (Hawkins & Roberts 2003). The stoplight parrotfish was one of the most abundant species in surveys and is usually the most frequently targeted by fishermen to serve in restaurants, however the population appears to be highly resilient, with impacts of fishing pressure rather being illustrated by its remarkable decrease in mean size. Overall, these results are consistent with other findings in the Caribbean region were parrotfish assemblages dominated by small sizes prevail as a consequence of overfishing (Hawkins & Roberts 2004).

The scarcity of large herbivore species has important effects in the ecosystem functionality because of their key role in reef dynamics (Bellwood et al. 2004). Recent studies have shown that herbivorous action varies according to parrotfish species, as they feed on different algae groups (Burkepile & Hay 2010). Burkepile & Hay (2010) suggest that this causes species-specific effects in the ecosystem, depending on the developmental state of the community, and provide evidence in support of protecting herbivore diversity, as this is crucial to promoting recovery processes that enhance overall resilience. In addition, small parrotfish seem to be less effective in their role as herbivores and bioeroders than their bigger counterparts. Lokrantz et al. (2008) showed that a reduction in body size of a parrotfish population causes disproportionate loss in their function. They revealed that to compensate for the loss of a single 35 cm individual of *Chlorurus sordidus*, 75 individuals of 15 cm are needed, suggesting that reefs with high parrotfish abundance may have functionality impairments if dominated by small sizes (Lokrantz et al. 2008).

Parrotfish fisheries in Colombia are not as common as in other regions (i.e., Indo-pacific) because traditionally these fish have not been considered commercial. Given that restaurants and fishermen trick tourists by taking advantage of their inability to identify parrotfish on their plates, the existence of a parrotfish market is a very delicate topic in CRSB and individuals can sometimes be very defensive about the subject. We believe it is for this reason that conflicting results were seen between fishermen surveys and group workshops in terms of parrotfish capture for marketing, however the following facts emerged: In 2000 parrotfish were first used to substitute snapper in tourist menus; parrotfish are captured mainly by pole spear (selective fishing gear targeting big individuals); parrotfish have low market value for fishermen but restaurants profit by selling them as snapper; and in day-tours (the most common tourism to CRSB) tourists pay a package that includes a fish lunch usually of “*mojarra lora*’’ or “*pargo risa*” (parrot mojarra and smiley snapper). Tourism to the CRSB has increased dramatically in recent years, causing a significant rise in commercial parrotfish fishing where local restaurant owners estimate current levels to be as high as 720-1800 parrotfish per day in high season in places such as Múcura Island. This kind of pressure in a parrotfish population assemblage already skewed to small sizes and low diversity is extremely worrying, moreover, other studies have shown that coral reef health in CRSB is already impacted, showing high algae abundance and low coral cover (Camargo et al. 2009).

There is a major lack of awareness about parrotfish by tourists and locals (from the city and local communities). Most tourists had never heard about parrotfish before. However, after learning the important role of parrotfish, most of them were inclined to support its protection. Likewise, during our awareness raising activities we noticed fishermen held misconceptions about the ecological role and feeding behaviour of parrotfish. Nevertheless, our activities targeting fishers seem to be effective in changing fishermen perspectives, and some of them were very surprised to find out about parrotfish importance and recognized the urgency of protection. Others went even further, by committing to stop its capture and showed initiative in spreading the information.

Awareness raising activities were very successful in delivering the conservation message and we believe the success relied heavily on building relationships within the community, showing respect, acceptance and interest in local culture, values and livelihoods. In general we found that these communities were committed to all the activities, providing us with their interest and support. Most importantly, they seemed to embrace the information showing clear intention during the entire project in conserving parrotfish and in starting dialogues with the environmental authorities to make participatory fishery agreements. However, in order to move forward they will need to resolve current tensions and conflict with environmental authorities, perhaps approached by both parties in establishing an appropriate forum.

Aswani & Sabetian (2010) studied the effects of urbanization on artisanal parrotfish fisheries in the Solomon Islands and found that in less than a year abundance dropped 50% in heavily fished areas, with medium and large individuals declining more sharply. The authors state that in communities with weak forms of customary management adjacent to urbanized areas, parrotfish can suffer a rapid decline in a short period of time (Aswani & Sabetian 2010). This scenario reveals that it is imperative to start accepting the formal existence of a parrotfish fishery in Colombia, since increasing tourism development and urbanization are bound to significantly stress artisanal fisheries, causing important cascading effects in coral reefs that need to be studied. Fishing is the main economic activity in these low-income communities, indicating a high level of dependence on marine resources, as principal source of income and protein. Moving towards sustainability will involve creating new economic alternatives. In addition, efforts should target not only fishermen who belong to the supply side of this market, but should also be directed to those who are buying the product. Our results suggest parrotfish should be prioritised in the environmental agenda of Colombia to increase awareness at a national level and start its protection. Parrotfish has potential for its use as a charismatic species or mascot for promoting the protection of coral reefs.

## 5. Conclusion

Parrotfish are important agents increasing resilience in coral reefs because of their role as bioeroders, herbivores and dispersers of zooxanthellae (Castro-Sanguino & Sanchez 2012). These ecological functions make these species a key conservation target to maintain and enhance reef health. In the CRSB, parrotfish populations are critical due to the degraded state of the reef. This evidence highlights that parrotfish should be put urgently in the environmental agenda.

We recommend the design and implementation of a Fisheries Plan of Action jointly between the fishing communities and the environmental authorities (Unidad de Parques Nacionales, INCODER, Coastal Guard). Such a plan should establish sustainable practice through regulation of fishing grounds, gear, minimum catch sizes and quota for particular species, including parrotfish. In order to make this fishery strategy work, cooperative alliances between different organizations must be consolidated (Coast Guard, MPA, INCODER, Capitanía de Puertos) and an effective surveillance and enforcement plan should be implemented.

Additionally, we also suggest that university and NGO research should be encouraged and their findings considered as baseline information in decision-making process; educational campaigns with tourists should be permanently established; and an environmental education plan should be integrated into the academic curriculum of the schools of Isla Grande, Múcura and El Islote.

## Bibliography

AGRRA. 2018. Atlantic and Gulf Rapid Reef Assessment (AGRRA): An Online Database of AGRRA coral reef survey data. Available: http://agrra.org

Aswani, S., and A. Sabetian. 2010. Implications of Urbanization for Artisanal Parrotfish Fisheries in the Western Solomon Islands. Conservation Biology 24:520–530.

Bejarano AC, Toline CA, Horsman JL, Zarza-González E, Cogollo K (2016) A climate change vulnerability framework for Corales del Rosario y San Bernardo National Natural Park, Colombia. Clim Res 70:1–18

Bellwood, D., T. Hughes, M. Nyström, and C. Folke. 2004. Confronting the coral reef crisis. Nature 429:827–833.

Burkepile, D. E., and M. E. Hay. 2010. Impact of Herbivore Identity on Algal Succession and Coral Growth on a Caribbean Reef. PLoS ONE 5:e8963.

Camargo, C., J. H. Maldonado, E. Alvarado, R. Moreno-Sánchez, S. Mendoza, N. Manrique, A. Mogollón, J. D. Osorio, A. Grajales, and J.A. Sánchez. 2009. Community involvement in management for maintaining coral reef resilience and biodiversity in southern Caribbean marine protected areas. Biodiversity and Conservation 18:935–956.

Castro-Sanguino, C., and J. A. Sanchez. 2012. Dispersal of Symbiodinium by the stoplight parrotfish Sparisoma viride. Biology Letters 8:282–286.

Durán, C.A. 2006. ¿Es nuestra isla para dos? Conflictos por el desarrollo y la conservación en Islas del Rosario, Cartagena. Tesis de grado Maestría en Antropología Social. Universidad de los Andes.

Durán, A., and R. Claro. 2008. Actividad alimentaria de los peces herbívoros y su impacto en arrecifes con diferente nivel de degradación antrópica. Revista de Biología Tropical 57:687–697.

Durán, C. A. 2009. Gobernanza en los Parques Nacionales Naturales colombianos. Revista de Estudios Sociales:60–73.

Fabricius, K. 2005. Effects of terrestrial runoff on the ecology of corals and coral reefs: review and synthesis. Marine Pollution Bulletin 50:125–146.

Gardner TA, Cote IM, Gill JA, Grant A, Watkinson AR. 2003. Long-term region-wide declines in Caribbean corals. Science 301: 958–960

Hawkins, J. P., and C. M. Roberts. 2004. Effects of artisanal fishing on Caribbean coral reefs. Conservation Biology 18:215–226. Wiley Online Library.

Hawkins, J., and C. Roberts. 2003. Effects of fishing on sex-changing Caribbean parrotfishes. Biological Conservation 115:213–226.

Hughes, T. P., M. J. Rodrigues, D. R. Bellwood, D. Ceccarelli, O. Hoegh-Guldberg, L. McCook, N. Moltschaniwskyj, M. S. Pratchett, R. S. Steneck, and B. Willis. 2007. Phase Shifts, Herbivory, and the Resilience of Coral Reefs to Climate Change. Current Biology 17:360–365.

Ledlie, M. H., N. A. J. Graham, J. C. Bythell, S. K. Wilson, S. Jennings, N. V. C. Polunin, and J. Hardcastle. 2007. Phase shifts and the role of herbivory in the resilience of coral reefs. Coral Reefs 26:641– 653.

Lokrantz, J., M. Nyström, M. Thyresson, C. Johansson, and undefined author. 2008. The non-linear relationship between body size and function in parrotfishes. Coral Reefs 27:967–974.

Martínez-Viloria, H. L. Martínez-Whisgman, A. Vargas-Pineda, J. Narváez Barandica. 2011. Efectos de la pesca sobre los recursos hidrobiológicos del Parque Nacional Natural Corales del Rosario y de San Bernardo. In: El Entorno Ambiental del Parque Nacional Natural Corales del Rosario y de San Bernardo. Eds: Zarza-González, E. Pp.273–289.

Morgan, K.M., S.K. Paul S. Kench. 2016. Parrotfish erosion underpins reef growth, sand talus development and island building in the Maldives. Sedimentary Geology. 341: 50–57.

Mumby, P. J., A. Hastings, and H. J. Edwards. 2007. Thresholds and the resilience of Caribbean coral reefs. Nature 450:98–101.

Pineda, I., L. Martinez, D. Bedoya, P. Caparroso, and J. Rojas. 2006. Plan De Manejo del Parque Nacional Natural Corales del Rosario y San Bernardo 2005-2009. Pages 1–306. Cartagena.

Pollnac, R., P. Christie, J. E. Cinner, T. Dalton, T. M. Daw, G. E. Forrester, N. A. J. Graham, and T. R. McClanahan. 2010. Marine reserves as linked social-ecological systems. PNAS 107:18262– 18265.

Shantz, A. A., Ladd, M. C., and Burkepile, D. E. 2020. Overfishing and the ecological impacts of extirpating large parrotfish from Caribbean coral reefs. Ecological Monographs 90(2):e01403. 10.1002/ecm.1403

